# The origin, divergence and inter-subspecies hybridization of house mice (*Mus musculus*)

**DOI:** 10.1101/2023.09.26.559459

**Authors:** Meidong Jing, Yingjie Chen, Wenbo Yuan, Ying Song, Xiaoxin Bi, Ling Huang

**Affiliations:** School of Life Sciences, Nantong University, Nantong, Jiangsu 226019, China; State Key Laboratory for Biology of Plant Diseases and Insect Pests, Institute of Plant Protection, Chinese Academy of Agricultural Sciences, Beijing 100193, China; Yantai Oceanic Environmental Monitoring Center, Ministry of Natural Resources, Yantai, Shandong 265503, China

## Abstract

House mice (*Mus musculus*) are the ancestors of laboratory mouse strains and an excellent model for evolutionary biology. The origin, divergence and inter-subspecies hybridization of the three main subspecies (*Mus musculus domesticus*, *M. m. musculus*, *M. m. castaneus*) were not well-resolved. Population genomic analyses with 349 samples from Eurasia confirmed the Himalayas-India-Pakistan junctional region as the origin centre. The divergence of *M. m. domesticus*, *M. m. musculus* and *M. m. castaneus* occurred ∼333.6, 308.3 and 134.4 thousand years ago, respectively, which located in different interglacial stages in Pleistocene. Hybridization among *M. m. domesticus*, *M. m. musculus*, and *M. m. castaneus* after secondary contact during spread and gene flow between the ancestral population and different subspecies produced distinct genomic mixtures in populations from different regions of Eurasia. Long-distance introgression from *M. m. musculus* into *M. m. castaneus* and from *M. m. castaneus* into *M. m. musculus* happened in southern China and northern China, respectively. And extensive genomic introgression occurred on both autosomes and the X chromosomes. Genomic introgression in hybrids from East Asia and Europe revealed that the level of genomic differentiation, recombination rate and benefits for the survival and reproduction of hybrids determined the introgression capability of distinct genomic regions in inter-subspecies hybridization of house mice. Unidirectional introgression of the Y chromosome made all wild mice in East Asia hold the *musculus*-type Y chromosome. Our study provided comprehensive insights into the origin, divergence and inter-subspecies’ genomic introgression of house mice.

## Introduction

House mice (*Mus musculus*), the ancestors of laboratory mouse strains and excellent models for evolutionary biology, which contain three main subspecies (*M. m. domesticus*, DOM; *M. m. musculus*, MUS; *Mus musculus castaneus*, CAS) occupying different regions of the world (Boursot et al. 1993, 1996; Bonhomme and Searle 2012; Phifer-Rixey and Nachman 2015; Ullrich and Tautz 2020). The origin centre of extent subspecies of house mice was still uncertain. Based on genetic variation in several molecular markers, Boursot and colleagues proposed that the northern part of the Indian subcontinent was the cradle of extant subspecies, and the westwards, northwards and southwards radiation of the ancestral population (AP) and subsequent genetic differentiation formed DOM, MUS and CAS, respectively (Boursot et al. 1993, 1996). However, some investigators believed that the Iranian plateau was also a candidate origin centre because this region had the highest diversity of lineages (Hardouin et al. 2015). The AP remains uncertain, and the genomic differentiation between the AP and various subspecies is unknown, so both origin theories lack more convincing evidence. It has been speculated that the divergence of house mice likely started ∼350-500 thousand years ago (kya) (Boursot et al. 1993; Geraldes et al. 2008, 2011), but the precise divergence time of each subspecies was also uncertain (Geraldes et al. 2008, 2011; Phifer-Rixey et al. 2020; Fujiwara et al. 2022).

Commensalism between humans and three subspecies of house mice evolved independently, which facilitated the passive colonization of DOM, MUS and CAS to different regions of the world (Boursot et al. 1993). Phylogeographic investigations have helped us to understand the regional migration histories of various subspecies (Rajabi-Maham et al. 2008; Searle et al. 2009; Jones et al. 2010; Bonhomme et al. 2011; Jing et al. 2014) and world-wide colonization routes (Bonhomme and Searle 2012; Phifer-Rixey and Nachman 2015; Harr et al. 2016). Different subspecies contacted secondarily during spread; then, a narrow hybrid zone between DOM and MUS formed in Europe, and hybrid zones between CAS and MUS formed in China and Japan (Yonekawa et al. 1988; Boursot et al. 1993; Duvaux et al. 2011; Bonhomme and Searle 2012; Jing et al. 2014; Phifer-Rixey and Nachman 2015; Harr et al. 2016). Such hybrid zones offer an excellent model for studying the mechanism of genomic introgression between closely related taxa in mammals and the genetic basis of reproductive isolation in speciation. Investigations of the hybrid zone in Europe revealed strong asymmetric introgression from DOM into MUS on autosomes (Payseur et al. 2004; Macholan et al. 2007; Geraldes et al. 2008; Teeter et al. 2008, 2010; Janoušek et al. 2015) and significantly reduced introgression on the X chromosome (Dod et al. 1993; Payseur et al. 2004; Macholan et al. 2007; Geraldes et al. 2008). X-Y or X-autosome incompatibility between the two subspecies (Campbell and Nachman 2014) has led to reduced fertility in male hybrids (Turner et al. 2012). However, hybrid zones in East Asia have rarely been explored (Yonekawa et al. 1988; Terashima et al. 2006; Jing et al. 2014), and the details on genomic introgression between CAS and MUS are unclear.

Here, 209 wild mice from China were resequenced to more than 10× depth on average. Combining available genome data of 140 wild mice from 19 Eurasian countries (Halligan et al. 2010; Harr et al. 2016; Fujiwara et al. 2022), our population genomic analyses confirmed that the Himalaya-India-Pakistan junctional region was the origin centre of house mice and that the divergence of DOM, MUS and CAS occurred in different interglacial stages of the Pleistocene. Genome-wide exploration of introgression in hybrids from East Asia and Europe revealed the determining factors that influence the introgression capability of distinct genomic regions. Our results provide a comprehensive picture of the origin, divergence and inter-subspecies hybridization of house mice.

## RESULTS

### Integration of genomic sequence data

We sequenced the whole genome of 209 wild mice from 32 localities of China, with coverage depths from 7.10× to 13.04×. Combining available genomic data (Halligan et al. 2010; Harr et al. 2016; Fujiwara et al. 2022), we compiled a collection of 349 wild mice (151 females and 198 males) from 19 countries (Fig. 1A; Supplemental Table S1). The genome sequence of *Mus spretus* from Spain was used as the outgroup. After mapping all sequence reads against the reference genome GRCm38.p6 and further filtering, we identified a total of 81,876,059 single-nucleotide polymorphisms (SNPs) on autosomes, 3,114,634 SNPs on the X chromosome and 10,874 SNPs on the nonrecombinant region of the Y chromosome (Supplemental Figs. S1, S2; Supplemental Table S2).

**Figure 1.**
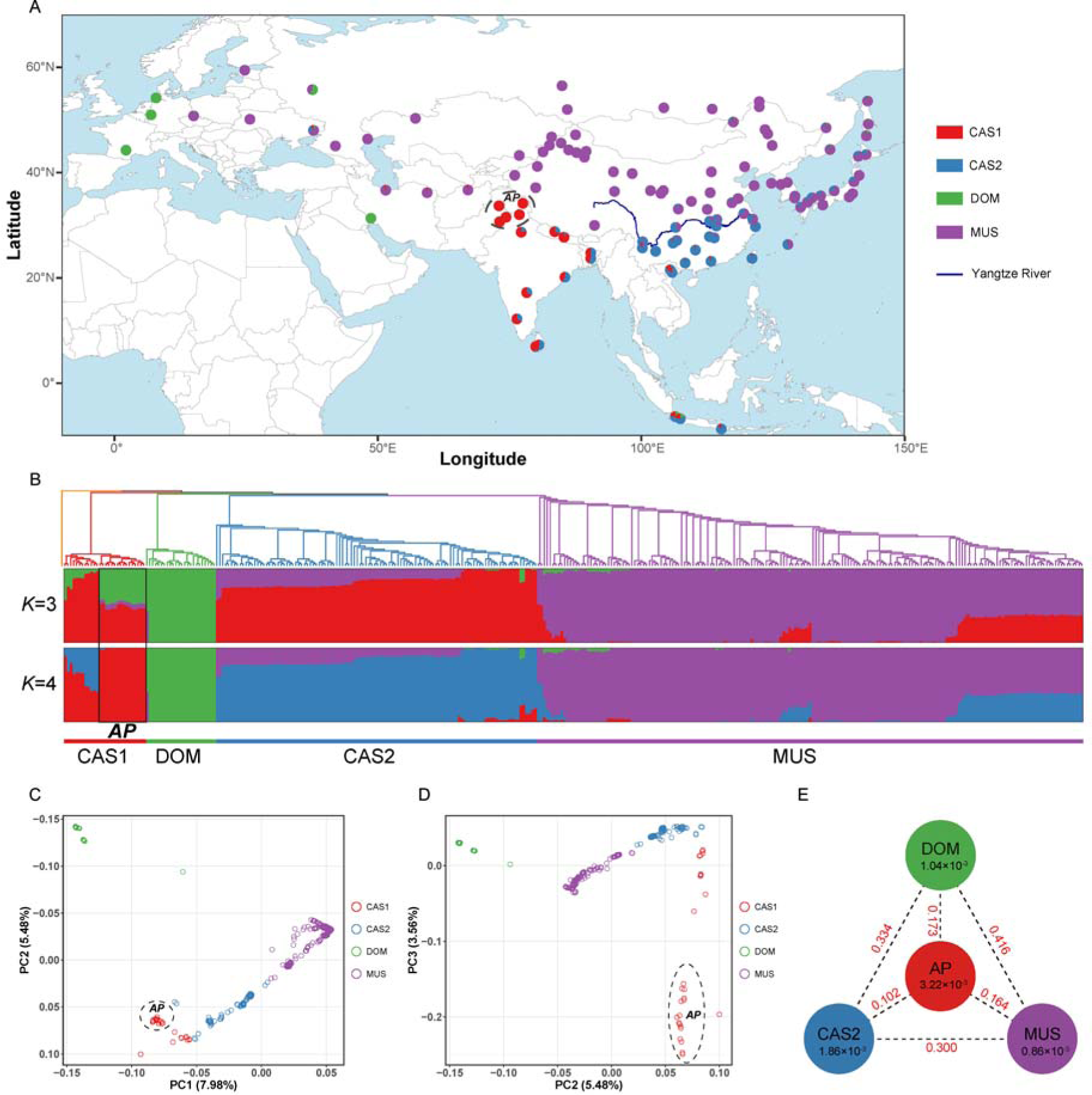
Population genetic analyses based on autosomal SNPs give clues on origin centre and genomic mixture between different clades of house mice. (A) Total of 349 samples from 112 localities (circles) of 19 Eurasian countries were involved in present study. The map was drawn using R ggplots2. The four colours in circles represented the ratio of the ancestral components inferred from ADMIXTURE analyses with *K* = 4. AP means the ancestral population, CAS means *M. m. castaneus*, MUS means *M. m. musculus*, DOM means *M. m. domesticus*. (B) The correspondence between neighbour-joining phylogenetic tree and ADMIXTURE analyses with *K* = 3 and 4. In NJ tree, the 349 samples were clustered into four distinct clades, which corresponded to the ancestral components inferred from ADMIXTURE analyses with *K* = 4. Sixteen individuals from the Himalayas-India-Pakistan junctional region were representatives of CAS1 in ADMIXTURE analyses, and they formed a single subclade of CAS1 in NJ tree. When *K* = 3, CAS1 and CAS2 merged into one component (CAS), and the 16 individuals comprised all three genomic components (CAS/DOM/MUS). The ADMIXTURE analyses also suggested that many samples from different regions of Eurasia exhibit genomic mixture between different clades. (C) Principal component analysis (PCA) differentiated 349 samples into two clusters along principal component 1 (PC1) and PC2. Individuals of DOM grouped into one specific cluster, and samples of CAS1, CAS2 and MUS scattered along a wide range of one cline. (D) Individuals of CAS1 and CAS2 were better differentiated along PC2 and PC3. CAS1 scattered over a wide range along PC3, and samples from the Himalayas-India-Pakistan junctional region were more divergent from CAS2. The population structure analyses based on the X chromosomal SNPs were shown in Supplemental Fig. S3. (E) Nucleotide diversity (π) and fixation index (*F_ST_*) across four clades of house mice. Value in each circle is the mean value of nucleotide diversity for each clade, and values in red on each line indicate pairwise population divergence between clades.

### Population genetic analyses give clues on origin centre and complex gene flow between different clades

We constructed the population structure including all 349 samples using autosomal and X-chromosomal SNPs. The neighbour-joining (NJ) trees showed that all samples were grouped into four distinct clades instead of three clades (Fig. 1B; Supplemental Fig. S3A). The CAS samples from the Indian subcontinent formed an independent clade (CAS1), and samples from Southeast Asia and China formed another clade (CAS2). In CAS1, 16 individuals from the junctional region of the southern foot of the Himalayas, northern India and Pakistan (IN1-10, 15-16, and PA1-4) were clustered into a subclade, and samples from other regions of the Indian subcontinent (Nepal, Bangladesh, Sri Lanka, hinterlands of India) were grouped into another subclade (Fig. 1B; Supplemental Fig. S3A).

We investigated the genetic structure for clusters (*K*) from 1 to 6 (Supplemental Fig. S4A). Analyses using autosomal SNPs obtained the lowest cross-validation error at *K*=4 (Supplemental Fig. S4b), and the four ancestral genomic components generally corresponded to CAS1, CAS2, DOM and MUS (Fig. 1B). Sixteen individuals from the Himalayas-India-Pakistan junctional region were representatives of CAS1; the samples from France and Germany and some individuals from Iran were representatives of DOM; the samples from Kazakhstan and Xinjiang of China were representatives of MUS; and other samples presented different levels of genomic mixture (Fig. 1A-B). At *K*=3, CAS1 and CAS2 merged into CAS. Individuals from the Himalayas-India-Pakistan junctional region comprised all three genomic components (CAS/DOM/MUS), and the samples from other regions of the Indian subcontinent, Southeast Asia and the south coast of China contained components of CAS and DOM, while most individuals from southern China presented genomic mixture of CAS and MUS (Fig. 1B). Analyses using X-chromosomal SNPs produced similar results, but the genomic mixture was reduced (Supplemental Fig. S3A). The residual fit from the maximum-likelihood tree deduced by TreeMix also indicated gene flow between CAS1 and CAS2, CAS2 and MUS, and MUS and DOM (Supplemental Fig. S5).

Principal component analysis (PCA) using autosomal SNPs differentiated all samples into two clusters along principal component 1 (PC1, 7.98% of variance) and PC2 (5.48% of variance) (Fig. 1C). Individuals of DOM (except RU9) clustered tightly, and other samples were scattered along a wide range of the CAS1/CAS2/MUS cline. CAS1 and CAS2 were better differentiated along PC2 and PC3 (3.56% of variance) (Fig. 1D). CAS1 was scattered over a wide range along PC3, and samples from the Himalayas-India-Pakistan junctional region were more divergent from CAS2. PCA analyses using X-chromosomal SNPs yielded similar results (Supplemental Fig. S3B-C).

We further applied the *D*-statistic to test the possibility of gene flow between different clades. The results strongly supported the hybridization between CAS2 and MUS in East Asia and between MUS and DOM in Europe (Fig. 3A; Supplemental Figs. S6-S9; Supplemental Tables S3-S5), as well as extensive gene flow between CAS1 and CAS2 in the Indian subcontinent (Supplemental Fig. S10; Supplemental Table S6). However, genomic introgression between MUS and CAS1 in Afghanistan and between CAS1 and CAS2 in Southeast Asia and China was limited (Supplemental Figs. S11, S12; Supplemental Tables S7, S8).

The analyses described above clearly showed that CAS2 in Southeast Asia and China diverged from CAS1 in the Indian subcontinent. Individuals from the Himalayas-India-Pakistan junctional region represent the AP of house mice, as they are representatives of CAS1 at *K*=4 and they comprise all three genomic components (DOM/MUS/CAS) at *K*=3 in ADMIXTURE analyses, and populations around the origin centre exhibit genomic mixture between the AP and different subspecies (Fig. 1A-B). Hybridization among DOM, MUS and CAS2 after secondary contact during spread and gene flow between the AP and different subspecies produced distinct genomic mixtures in different geographic populations of Eurasia (Fig. 1A), which could result in the discordant topology of NJ trees inferred from autosomal SNPs and X-chromosomal SNPs (Fig. 1B; Supplemental Fig. S3A).

To eliminate the impact of introgression on phylogeny, we selected ten representatives with |*D|* < 0.05 and |Z| score < 3 in *D*-statistics for each clade (Supplemental Figs. S6-S14; Supplemental Tables S1, S3-S10). The population structure inferred from the 40 representatives verified the validity of these representatives (Supplemental Fig. 15A-C). We then estimated the nucleotide diversity (π) of four clades and genomic differentiation (*F*_ST_) between different clades. The nucleotide diversity for each clade was 3.22×10^−3^ (AP), 1.86×10^−3^ (CAS2), 1.04×10^−3^ (DOM) and 0.86×10^−3^ (MUS), respectively (Fig. 1E; Supplemental Figs. S16A, S17). The linkage disequilibrium decayed more rapidly in clades with higher levels of diversity (Supplemental Fig. S16B). The level of genomic differentiation between the AP and the three subspecies (AP/CAS2: 0.102, AP/MUS: 0.164, AP/DOM: 0.173) were evidently lower than those between the three subspecies (CAS2/MUS: 0.300, CAS2/DOM: 0.334, DOM/MUS: 0.416) (Fig.1E; Supplemental Fig. S18). For all three subspecies, with the increase in geographic distance from the AP, nucleotide diversity generally decreased, whereas genomic differentiation with the AP was increased. The nucleotide diversity of different geographic populations were negatively associated with the genomic differentiation between the AP and these populations (Pearson’s coefficient: *R* = −0.98 with *P* < 2.2 × 10^−16^) (Supplemental Fig. S16C). These results further confirm that the Himalayas-India-Pakistan junctional region is the cradle of house mice (Supplemental Fig. S19).

### Subspecies’ divergence of house mice occurred during different interglacial stages

We applied the *f_2_* and *f_3_* statistics to explore the genetic relationship among the four clades of house mice, and the results supported the closest relationship between the AP and CAS2 and the distant relationship between DOM and other clades (Supplemental Fig. 20). So, the results of phylogenetic tree including 40 representatives (Supplemental Fig. 15A), genomic differentiation among different clades (Fig. 1E; Supplemental Fig. S18) and *f_3_* statistics (Supplemental Fig. S20) together validate the earliest divergence of DOM and the most recent divergence of CAS2 from the AP.

To estimate the divergence times of the three subspecies, we selected six representatives with low rates of missing data for each clade (Supplemental Table S1). Analyses using the joint site frequency spectrum of 24 representatives (Supplemental Fig. S21) revealed that DOM, MUS and CAS2 diverged from the AP ∼333.6 kya (CI: 321.5-345.6), ∼308.3 kya (CI: 297.1-320.0) and ∼134.4 kya (CI: 129.1-140.6), respectively (Fig. 2A; Supplemental Fig. S22). The divergence of DOM and MUS occurred in the Mindel-Riss interglaciation in Europe and interglacial stages in the Penultimate Glaciation in China, and the divergence of CAS2 occurred in the interglaciation before the last glacial period (Yi et al. 2007).

**Figure 2.**
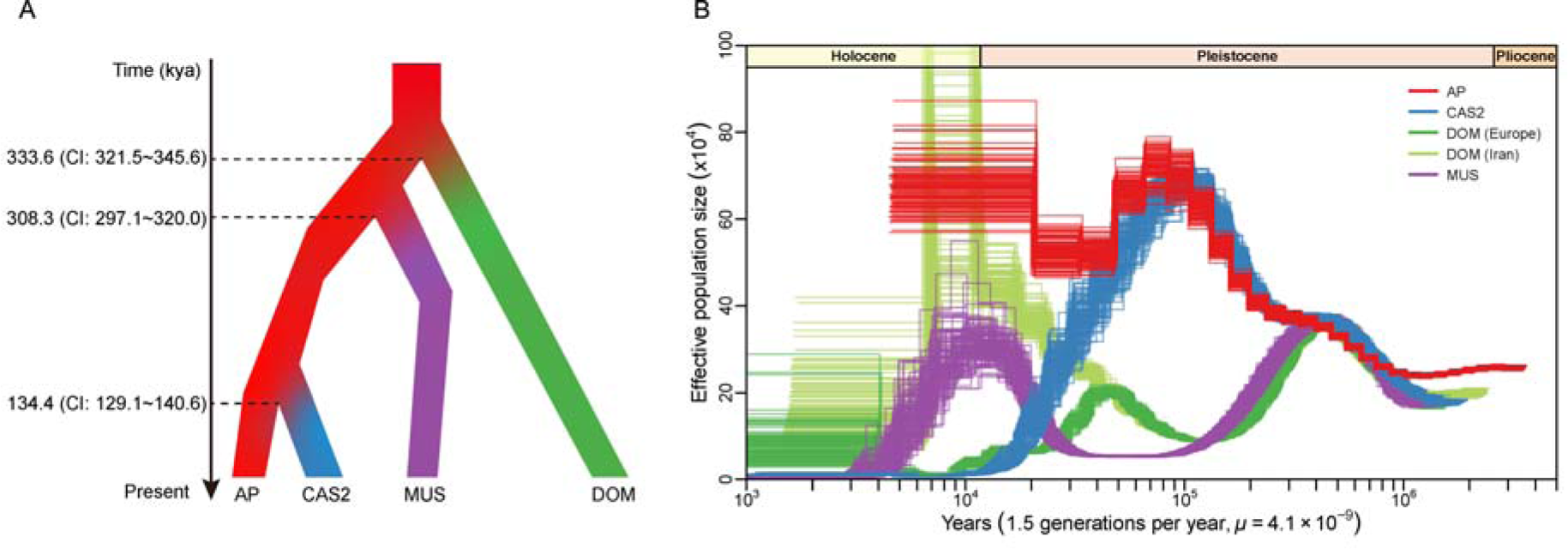
Divergence and demographic inference of house mice. (A) Divergence times of *M. m. domesticus*, *M. m. musculus* and *M. m. castaneus* were estimated using fastsimcoal2. Point estimates and the 95% confidence interval (CI) are shown on the left. (B) The past demographic patterns of the four clades using the pairwise sequentially Markovian coalescent. The trajectories of *M. m. domesticus* and *M. m. musculus* separated sharply from those of the ancestral population ∼300-350 kya. Five individuals representing the ancestral population, *M. m. domesticus*, *M. m. musculus* and *M. m. castaneus* were included in the analyses, with 100 bootstrap replicates. The demographic trajectories of *M. m. domesticus* from Iran and Europe were different, so representative from both regions were used in the analyses.

We deduced the past demographic patterns of the four clades applying the pairwise sequentially Markovian coalescent method using representatives with sequence coverage depth ≥ 18× (Supplemental Table S1). The results showed that trajectories of individuals from the same population were similar (Supplemental Fig. S23), and that each clade had a specific demographic pattern (Fig. 2B). The DOM and MUS trajectories separated sharply from those of the AP ∼300-350 kya (Fig. 2B), which was coincident with the divergence times of DOM and MUS (Fig. 2A; Supplemental Fig. S22).

### Extensive introgression between CAS2 and MUS occurred on both autosomes and the X chromosome

The population structure inferred from both autosomal SNPs and X-chromosomal SNPs indicated extensive introgression between CAS2 and MUS in China and Japan (Fig. 1B-D; Supplemental Figs. S3A-C, S5). In the *D*-suite analyses, five populations in China (WH, XG, YM, PX, and GZ, n≥5) presented signals for introgression from MUS into CAS2, while three populations in China (SH, XZ and NY, n≥5) and all samples from Japan (JP, n=18) showed signals for introgression from CAS2 into MUS (Supplemental Figs. S6, S24). The *D*-statistics for individuals revealed 107 hybrids in MUS from 20 localities and 90 hybrids in CAS2 from 12 localities of China (Fig. 3A; Supplemental Figs. S7, S8; Supplemental Tables S3, S4), which suggest long-distance introgression between the two subspecies starting along the sides of the Yangtze River in China.

**Figure 3.**
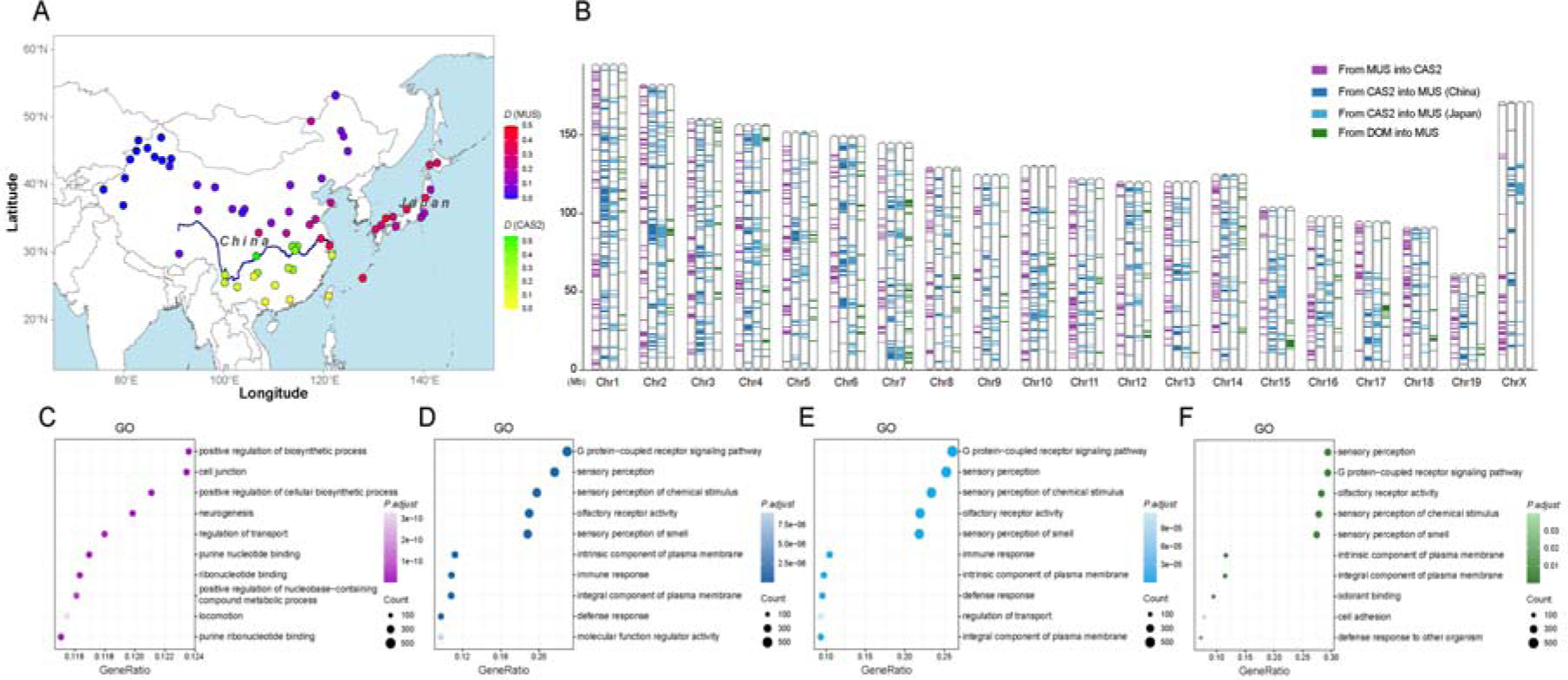
Genomic introgression between different subspecies of house mice. (A) The *D* values of different geographic populations in China and Japan. Long distance genomic introgression between *M. m. musculus* and *M. m. castaneus*, starting along the sides of the Yangtze River, had occurred in China. (B) Genomic distributions of introgressive segments from *M. m. musculus* into *M. m. castaneus* in China (purple), from *M. m. castaneus* into *M. m. musculus* in China (deep blue), from *M. m. castaneus* into *M. m. musculus* in Japan (light blue), and from *M. m. domesticus* into *M. m. musculus* in the Czech Republic (green). Most introgressive segments from *M. m. musculus* into *M. m. castaneus* and from *M. m. castaneus* into *M. m. musculus* were not overlapped, while introgression from *M. m. castaneus* into *M. m. musculus* that happened independently in China and Japan shared near half number of introgressive segments. Extensive introgression on the X chromosome occurred between *M. m. castaneus* and *M. m. musculus*, while no evident introgression on the X chromosome happened between *M. m. domesticus* and *M. m. musculus*. Functional enrichment of introgressive genes from *M. m. musculus* into *M. m. castaneus* in China (C) from *M. m. castaneus* into *M. m. musculus* in China (D), from *M. m. castaneus* into *M. m. musculus* in Japan (E), and from *M. m. domesticus* into *M. m. musculus* in the Czech Republic (F). The top 10 GO annotations with adjusted *P* < 0.05 are shown. The results suggested that genes transferred into *M. m. castaneus* and genes transferred into *M. m. musculus* show distinct bias on biological functions.

In the subsequent analyses, we clustered hybrids with introgression from MUS into CAS2 in China, hybrids with introgression from CAS2 into MUS in China and hybrids from Japan into distinct hybrid group, respectively. We used the top 5% of modified *f*-statistic (*f*_dM_) values as the criterion to screen the introgressive segments on autosomes and the X chromosome in three hybrid groups, because it is difficult to screen the introgressive windows with *f*-statistics (*f*_d_) in hybrid populations with high level of introgression. We identified 854 autosomal and 41 X-chromosomal introgressive segments (introgression from MUS into CAS2), 950 autosomal and 40 X-chromosomal introgressive segments (introgression from CAS2 into MUS in China) and 791 autosomal & 22 X-chromosomal introgressive segments (introgression from CAS2 into MUS in Japan), respectively (Fig. 3B; Supplemental Figs. S25, S26; Supplemental Tables S11-S14). The total length of introgressive segments was ∼8% of the autosomal genome and was ∼6% - 7% of the X chromosome. Notably, introgression from CAS2 into MUS in China and Japan shared 411 introgressive segments (2,223 windows), and the total length was ∼3.29% of the autosomal genome (Fig. 3B; Supplemental Fig. S27A; Supplemental Table S15).

We estimated the local recombination rates along the genomes of CAS2 and MUS (Supplemental Fig. S28), and calculated the expected shared ancestral segments and probabilities of retaining long ancestral segments (Supplemental Table S16). The results suggested that introgressive segments identified in the two genomes were unlikely remnants of incomplete lineage sorting. Introgressive segments identified in the two genomes were generally located in regions with comparative higher recombination rates and lower levels of genomic differentiation (Supplemental Fig. S28). For introgressive windows, the *f*_dM_ values were negatively associated with the genomic differentiation between the two subspecies (Pearson’s coefficients: *R* = −0.27 with *P* < 2.2 × 10^−16^ in introgression from MUS into CAS2; *R* = −0.26 with *P* < 2.2 × 10^−16^ in introgression from CAS2 into MUS), and were positively associated with the local recombination rates (Pearson’s coefficients: *R* = 0.16 with *P* = 1.3 × 10^−8^ in introgression from MUS into CAS2; *R* = 0.11 with *P* = 2.4 × 10^−4^ in introgression from CAS2 into MUS) (Supplemental Fig. S29A-B).

For hybrids from China, most of the genomic admixtures occurred at approximately 100 to 9,000 generations (Supplemental Tables S17, S18). Assuming an average reproductive rate of 1.5 generations per year (Phifer-Rixey et al. 2020), the hybridization between CAS2 and MUS started ∼6 kya, which is coincident with the development of ancient agriculture (Jones and Liu 2009) and the population expansion of MUS and CAS2 in China (Jing et al. 2014).

### Genes in introgression from MUS into CAS2 and from CAS2 into MUS shew distinct bias on biological functions

In introgressive segments on autosomes, we identified 1,704 genes (introgression from MUS into CAS2 in China), 2,207 genes and 2,254 genes (introgression from CAS2 into MUS in China and Japan), respectively (Supplemental Tables S11-S13). Introgression from CAS2 into MUS in China and Japan shared 1,049 genes (Supplemental Table S15).

Gene Ontology (GO) analyses revealed that a portion of the introgressive genes function on the plasma membrane, with more genes functioning in the cytoplasm and nucleus (Supplemental Table S19). GO and Kyoto Encyclopedia of Genes and Genomes (KEGG) analyses showed that introgressive genes from MUS into CAS2 significantly enriched in biological functions related to regulation of metabolism, reproduction and development (especially neurogenesis), cell transportation, cell death and immune response (Fig. 3C, Supplemental Fig. 30A; Supplemental Tables S20, S21), while introgressive genes from CAS2 into MUS were predominantly concentrated in biological functions including sensory perception of chemical stimulus, immune response and cell transportation (Fig. 3D-E; Supplemental Figs. S27B-C, S30B-C; Supplemental Tables S22-S27).

In introgressive segments on the X chromosome, we found 54 genes (introgression from MUS into CAS2), 39 genes and 29 genes (introgression from CAS2 into MUS in China and Japan), respectively. According to gene annotation in NCBI, 19 of these genes are specifically expressed in adult testes and related to male reproduction (Supplemental Table S14). The extensive introgression on the X chromosome indicated that the X-Y and X-autosome incompatibilities between CAS2 and MUS are weaker than those between DOM and MUS.

### Introgression between MUS and CAS2 are beneficial for individual survival and reproduction of hybrids

To explore if the introgression between CAS2 and MUS are beneficial for adaptation of hybrids, we screened overlapped genomic regions locating in the top 5% of *f*_dM_, *F*_ST_ (backcrossed parents/hybrids) and π ratio (backcrossed parents/hybrids). We found that 371 introgressive genes from 184 segments had been fixed in hybrids with introgression from MUS into CAS2 (Supplemental Fig. S31A-D; Supplemental Table S28). These genes significantly enriched in biological functions related to regulation of reproduction and development (especially neurogenesis), metabolism, cell transportation and immune response (Supplemental Fig. S32A-B; Supplemental Tables S29, S30). Genes playing important roles in functional maintenance of the nervous system, visual development, production of cerebrospinal fluid, and inner ear homeostasis (*Slc4a10* on chromosome 2) (Jacobs et al. 2008; Potter et al. 2016), spermatogenesis and fertilization of humans and mice (*Mfsd14a* on chromosome 3 and *Actl7a*/*Actl7b* on chromosome 4) (Doran et al. 2016; Xin et al. 2020) and the immune response (*Ccl3*, *4*, *6* and *Wfdc*17, 18, 21 on chromosome 11) presented strong signals of adaptive introgression (Fig. 4A-E).

**Figure 4.**
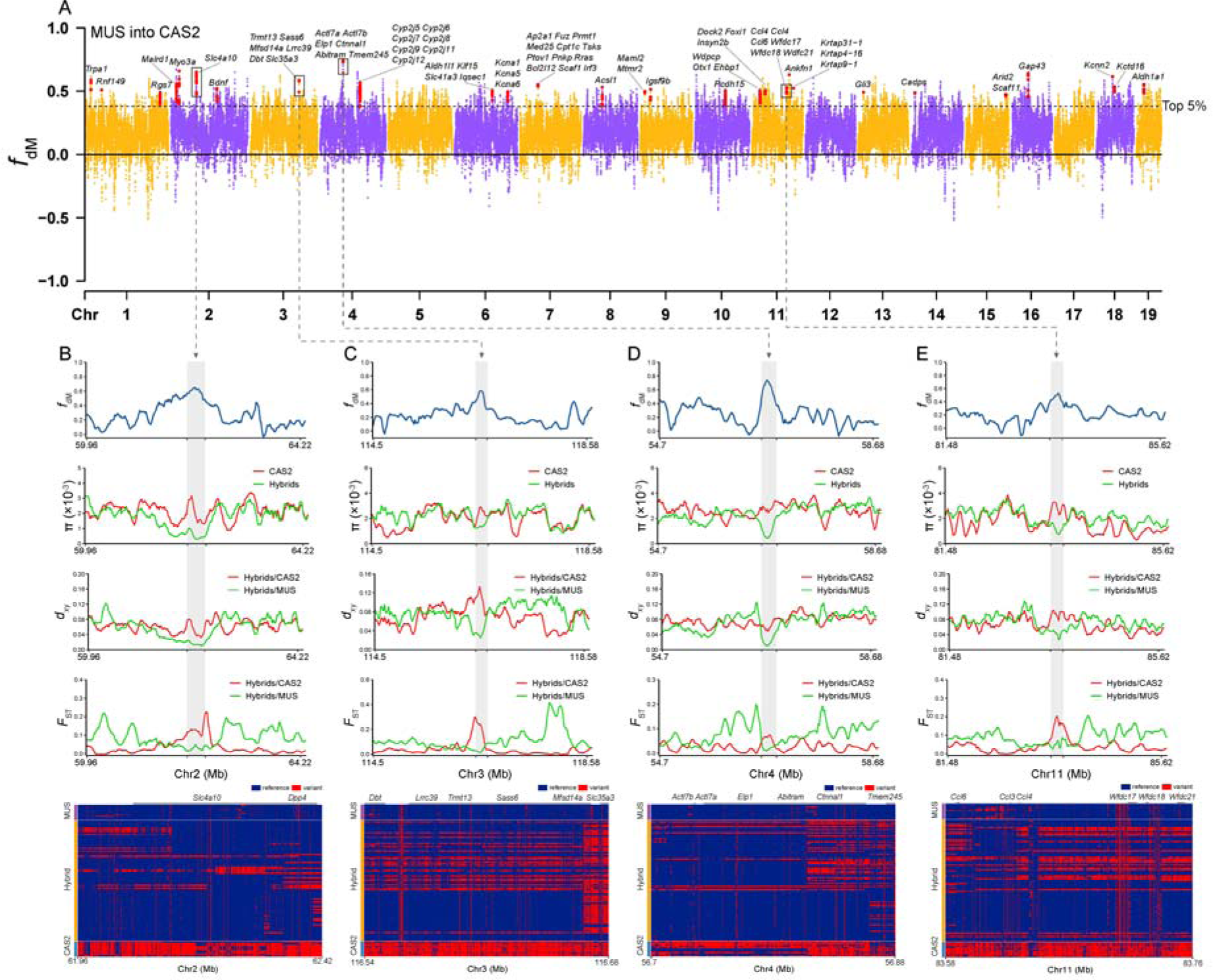
Adaptive introgression from *M. m. musculus* into *M. m. castaneus* in China. (A) Introgressive regions from *M. m. musculus* into *M. m. castaneus* identified in hybrids from China. A modified *f*-statistic (*f*_dM_) for 100-kb windows with 20 kb steps is plotted along the chromosomes. Each dot represents a 100-kb window. The black dotted horizontal line corresponds to the top 5% threshold. Some genes with strong signals of adaptive introgression are shown. Segments with strong signals of adaptive introgression on chromosome 2 (B), chromosome 3 (C), chromosome 4 (D) and chromosome 11 (E) contain genes that play great roles in functional maintenance of the nervous system (*Slc4a10* on chromosome 2), spermatogenesis and fertilization of mice (*Mfsd14a* on chromosome 3 and *Actl7a/Actl7b* on chromosome 4) and the immune response (*Ccl3*/*4*/*6* and *Wfdc17/18/21* on chromosome 11). The *f*_dM_ and nucleotide diversity (π) of hybrids and *M. m. castaneus* representatives, mean pairwise sequence divergence (*d*_xy_) and genomic differentiation (*F*_ST_) between hybrids and their parents, and haplotype patterns of hybrids and their parents compared to the reference genome GRCm38.p6 are shown for each region.

In introgression from CAS2 into MUS, we screened out 130 introgressive segments including 169 genes and 28 introgressive segments comprising 41 genes that had been fixed in hybrids from China and Japan, respectively (Supplemental Figs. S33, S34; Supplemental Tables S31, S32). These genes were predominantly concentrated in olfactory perception and immune response (Supplemental Figs. S35, S36; Supplemental Tables S33-S35).

The olfactory system of mice is responsible for recognizing various odours and pheromones to modulate individual social and reproductive behaviours (Del Punta et al. 2002). Vomeronasal and olfactory receptor genes on chromosomes 3, 7, 10, 13, 14 & 19 exhibited signals of adaptive introgression (Fig. 5A). Hybrids in China and Japan shared 14 genes with signals of adaptive introgression, including four genes that play important roles in retrovirus resistance (*Fv1*) (Yap et al. 2014), carcinogenesis (*Miip* and *Plod1*) (Chen et al. 2016; Wang et al. 2021) and spermatogenesis and meiosis (*Mfn2*) (Wang et al. 2022) on chromosome 4 and vomeronasal receptor genes (*Vmn2r51*-*52*) on chromosome 7 (Fig. 5A-C). Our results suggest that genomic regions with benefits for the survival and reproduction of hybrids are more prone to introgression, and a portion of introgressive genes have been fixed in hybrids by adaptative selection.

**Figure 5.**
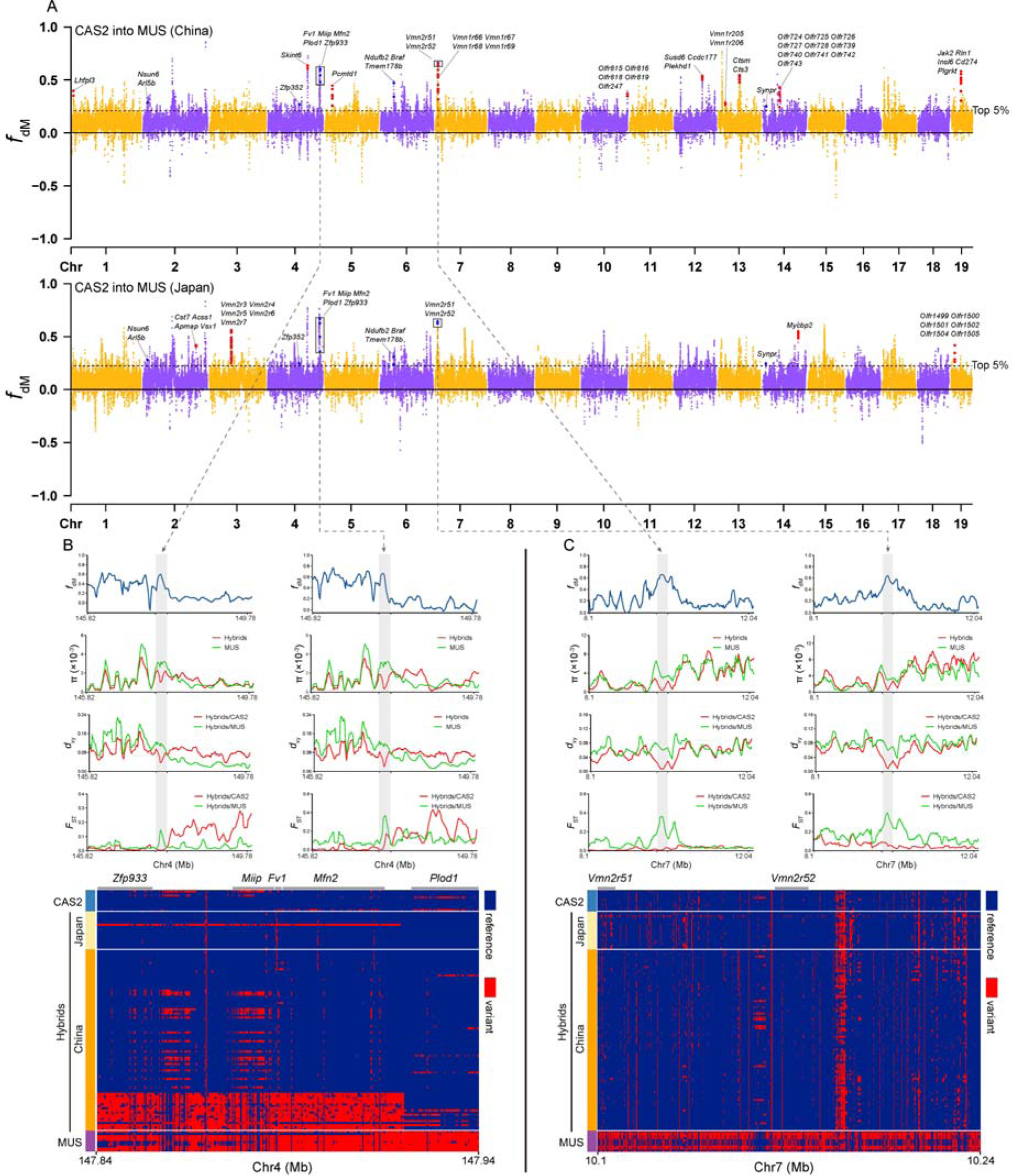
Adaptive introgression from *M. m. castaneus* into *M. m. musculus* in China and Japan. (A) Introgressive regions from *M. m. castaneus* into *M. m. musculus* identified in hybrids from China and Japan. A modified *f*-statistic (*f*_dM_) for 100-kb windows with 20 kb steps is plotted along the chromosomes. Each dot represents a 100-kb window. Red dots represent windows identified specifically in each group, while blue dots represent windows identified in both groups. The black dotted horizontal line corresponds to the top 5% threshold. Some genes with strong signals of adaptive introgression are shown. Adaptive introgression shared by hybrids in China and Japan on chromosome 4 (B) and chromosome 7 (C). The segment on chromosome 4 comprises four genes playing important roles in retrovirus resistance (*Fv1*), carcinogenesis (*Miip* and *Plod1*), spermatogenesis and meiosis (*Mfn2*), and the segment on chromosome 7 contains vomeronasal receptor genes (Vmn2r51-52). The *f*_dM_ and nucleotide diversity (π) of hybrids and *M. m. musculus* representatives, mean pairwise sequence divergence (*d*_xy_) and population differentiation (*F*_ST_) between hybrids and their parents, and haplotype patterns of hybrids and their parents compared to the reference genome GRCm38.p6 are shown.

### Unidirectional introgression on the Y chromosome between CAS2 and MUS

We constructed the phylogeny of 198 males using autosomal SNPs and Y-chromosomal SNPs, respectively. The trees presented interesting results: CAS1, DOM and MUS possess their own specific Y haplogroups, while CAS2 does not have a distinctive Y haplogroup (Fig. 6A). CAS2 individuals in Southeast Asia possess CAS1-type Y, while all CAS2 individuals in southern China hold MUS-type Y (Fig. 6A-B). The statistical parsimony network of Y-chromosomal haplotypes showed the same results (Supplemental Fig. S37). The geographic distribution of Y haplogroups (Fig. 6B) and migration routes of MUS and CAS2 in Asia (Supplemental Fig. S19) suggest unidirectional introgression of the MUS-type Y into CAS2 in China, which led to the replacement of the Y chromosome in CAS2 by the MUS-type Y, even in populations without evident introgression on autosomes and the X chromosome. The asymmetric introgression of MUS-type Y into DOM also occurred in some transects of the European hybrid zone (Munclinger et al. 2002; Jones et al. 2010). The ultra-strong introgression ability of MUS-type Y into other subspecies of house mice is an attractive topic for future studies.

**Figure 6.**
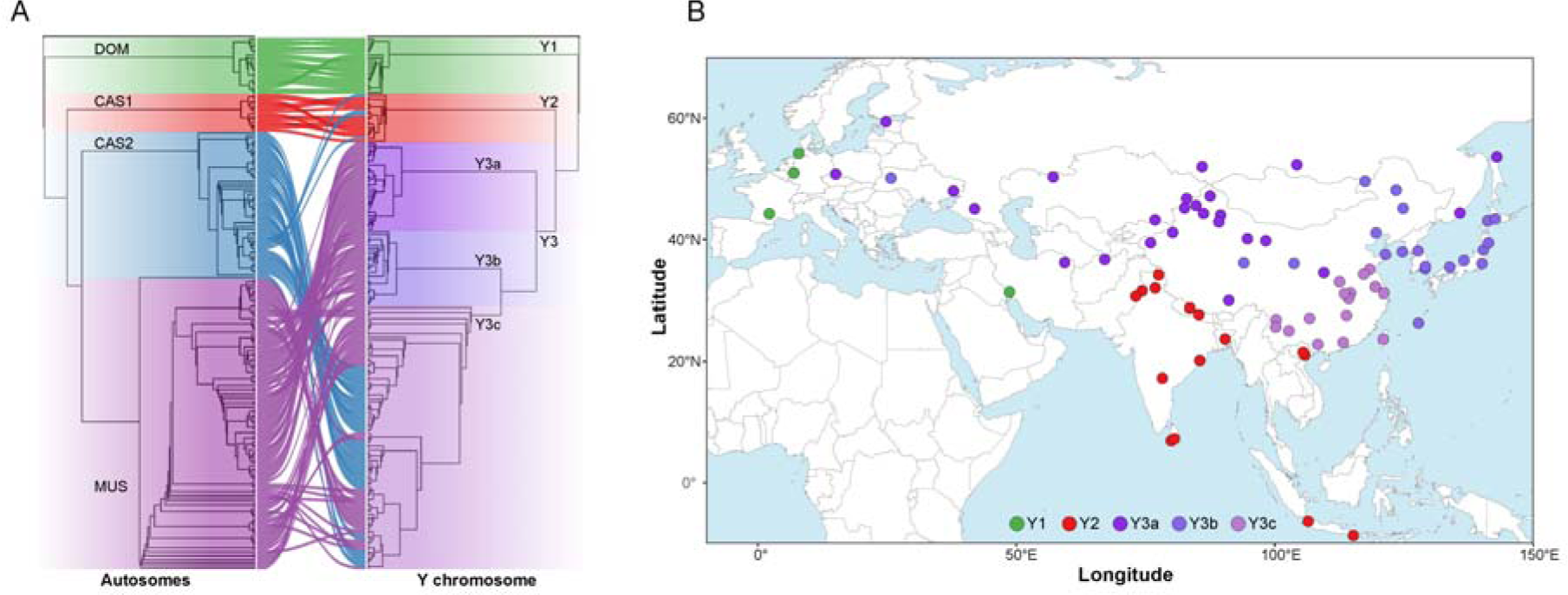
The paternal phylogeny of house mice revealed unidirectional introgression of the Y chromosome from *M. m. musculus* into *M. m. castaneus*. (A) The neighbour-joining phylogeny including 198 males inferred from autosomal SNPs (left) and Y-chromosomal SNPs (right). The populations from the Indian subcontinent (CAS1), *M. m. domesticus* and *M. m. musculus* possess their own specific Y haplogroups, while *M. m. castaneus* in Southeast Asia and East Asia (CAS2) do not have distinctive Y haplogroup. Individuals in Southeast Asia possess CAS1-type Y, while all *M. m. castaneus* individuals in southern China hold MUS-type Y. Graphs were drawn using the ape package. (B) The geographic distribution of Y-chromosome haplogroups in Eurasia. The colours in circles represented specific Y-chromosome haplogroups. The results suggest unidirectional introgression of the MUS-type Y chromosome into *M. m. castaneus* in China.

### Genomic introgression between DOM and MUS

Comparison on genomic introgression in hybrids from East Asia and Europe can help us to better understand the mechanism for inter-subspecies hybridization of house mice. In the genetic structure analyses and *D*-statistics with autosomal SNPs, we found six samples from the Czech Republic with evident signals for introgression from DOM into MUS (Fig. 1B; Supplemental Figs. S9, S38A; Supplemental Table S5). However, no evident introgression was identified based on the X-chromosomal SNPs (Supplemental Figs. S3A, S38B).

We screened the introgressive segments on autosomes in these six hybrids using a criterion of *f*_dM_ values locating in Benjamini-Hochberg false discovery rate (FDR) 5% significance level, and identified 342 segments containing 897 genes (Fig. 3B; Supplemental Fig. S39; Supplemental Table S36). The *f*_dM_ values for introgressive windows were also negatively associated with the genomic differentiation between DOM and MUS (Pearson’s coefficients: *R* = −0.12 with *P* = 2.4 × 10^−5^) and positively associated with the local recombination rate in MUS genome (Pearson’s coefficients: *R* = 0.078 with *P* = 7.2 × 10^−3^) (Supplemental Fig. S29C). The total length of introgressive segments was ∼3.1% of the autosomal genome. The introgressive genes (511 genes) functioning in the cytoplasm and nucleus were far more than those (241 genes) functioning on the plasma membrane (Supplemental Table S19). The introgressive genes significantly enriched in biological functions including sensory perception, immune response and cell transportation (Fig. 3F; Supplemental Fig. S40; Supplemental Tables S37, S38). We found that 51 introgressive genes from 15 segments had been fixed in hybrids, among which 27 genes encoded olfactory receptors (Supplemental Fig. S41; Supplemental Table S39). The introgression of genes related to olfactory receptors and immune response in the European hybrid zone was also reported in study with 1,316 autosomal SNPs (Janoušek et al. 2015).

## Discussion

The evolution history of house mice had attracted the interests of many researchers for several decades, but the origin centre and divergence times of three main subspecies (DOM, MUS and CAS2) were still uncertain (Boursot et al. 1993, 1996; Geraldes et al. 2008, 2011; Hardouin et al. 2015; Phifer-Rixey et al. 2020; Fujiwara et al. 2022). Boursot et al (1993) pointed out that the ancestral population of house mice was not attributable to any of subspecies though they were most likely CAS-like. However, for a long time, the ancestral population has not been ascertained, and all populations from the Indian subcontinent and Southeast Asian & East Asian were classified as *M. m. castaneus*, which brought difficulties to investigations on subspecies divergence (Geraldes et al. 2008, 2011; Phifer-Rixey et al. 2020; Fujiwara et al. 2022). The integration and addition of genomic data for wild mice from different regions of Eurasia, especially from the Indian subcontinent and Southeast Asian and East Asian, allow us to better distinguish the genomic differences between the ancestral population and new-born subspecies, then help us to define the geographic scope of the origin centre of house mice. Our population structure analyses with whole-genome data of 349 samples from 19 Eurasian countries revealed that the individuals from the Himalayas-India-Pakistan junctional region represent the ancestral population of house mice. *M. m. castaneus* in Southeast Asia & East Asia (CAS2) had diverged from the ancestral population, and populations in most regions of the Indian subcontinent exhibit genomic mixture between the ancestral population and CAS2. Our results confirm the origin centre of house mice with solid evidence from genomic data.

Our population structure analyses also showed that genomic mixture between different clades of house mice were more complexed than we had thought. Hybridization between different subspecies after secondary contact during world-wide colonization in recent history and ancient gene flow between the ancestral population and extant subspecies produced distinct genomic mixtures in populations from different regions of Eurasia. This is an important factor to explain the discordant phylogenies of house mice inferred from DNA markers of different genomic regions in present study and previous investigations (Lundrigan et al. 2002; Geraldes et al. 2008; Liu et al. 2008; Morgan and Pardo-Manuel de Villena 2017). After eliminating the impact of introgression, our phylogenetic relationship analyses further confirmed the earliest divergence of DOM and the most recent divergence of CAS2 from the ancestral population. With ascertain of the ancestral population, our estimation revealed that the divergence of DOM, MUS and CAS2 occurred during different interglacial stages in Pleistocene beginning ∼350 kya, which was also supported by deduction on the past demographic patterns of four clades. Our results were not only consistent with previous proposal that the divergence of house mice started ∼350-500 kya (Boursot et al. 1993; Geraldes et al. 2008, 2011) but also in accordance with the fact that warm and wet climates in interglacial stages facilitated population expansion and subsequent genetic differentiation (Hewitt 2000).

Hybrid zones between different subspecies of house mice is an ideal model to understand the mechanism of genomic introgression between closely related taxa in mammals under natural condition. The hybrid zone between DOM and MUS in Europe is narrow (Boursot et al. 1993; Duvaux et al. 2011; Bonhomme and Searle, 2012; Phifer-Rixey and Nachman, 2015; Harr et al. 2016), and the hybridization exhibits strong asymmetric introgression from DOM into MUS on autosomes (Payseur et al. 2004; Macholan et al. 2007; Geraldes et al. 2008; Teeter et al. 2008, 2010; Janoušek et al. 2015) and significantly reduced introgression on the X chromosome (Dod et al. 1993; Payseur et al. 2004; Macholan et al. 2007; Geraldes et al. 2008; present study). Our analyses on hybrid zones in China and Japan with whole-genome data revealed great differences on hybridization between MUS and CAS2 with that between MUS and DOM. Long-distance introgression from MUS into CAS2 and from CAS2 into MUS occurred in southern China and northern China, respectively, and 18 samples from different regions of Japan were all hybrids. Moreover, extensive introgression between MUS and CAS2 occurred on both autosomes and the X chromosome. Genetic incompatibility caused by genomic differentiation produce great resistance to gene flow between two different subspecies. The lower level of genomic differentiation between CAS2 and MUS (*F*_ST_: 0.300) than that between DOM and MUS (*F*_ST_: 0.416) was accordant with the evidently weaker genetic counteraction between MUS and CAS2.

We found that introgressive segments identified in introgression from MUS into CAS2, from CAS2 into MUS and from DOM into MUS are generally located in regions with comparatively lower levels of differentiation but higher recombination rates. And the *f*_dM_ values of introgressive windows were negatively associated with the genomic differentiation between the parental subspecies, and were positively associated with the local recombination rate in recipient genome. These results suggest that the genomic differentiation and the recombination rate are important factors influencing the introgression capability of distinct genomic regions in inter-subspecies hybridization of house mice. Genomic regions with higher recombination rates generally accumulate lower level of differentiation, and species barriers are more porous in genomic regions with higher recombination rates, which is also supported by recent investigations on introgression in both animals and plants (Martin et al. 2019; Christmas et al. 2021; Fu et al. 2022; Feng et al. 2023). The more interesting finding was that the genes transferred into MUS from both DOM and CAS2 are significantly enriched in similar biological functions, including olfactory perception, immune response and cell transportation, although most introgressive segments identified in hybrids from Europe and East Asia did not overlap. Obviously, this situation cannot be explained just by genomic differentiation and recombination rates. Further analyses showed that a portion of introgressive segments showed strong signals of adaptative selection, and that introgressive genes in these segments had been fixed in hybrids. For each analysed hybrid group, the genes in introgressive segments and adaptative introgression significantly concentrated in similar biological functions, which suggest that genomic regions that are beneficial for adaptation of hybrids are more prone to introgression. The olfactory sensory system and immune system are essential for the environmental adaptation and reproductive behaviour regulation of many taxa of animals, and the inter-species introgression of related genes have also been identified in several mammal taxa (Teng et al. 2017; Chen et al. 2018; Zheng et al. 2020; Chen et al. 2021). More than one fifth of introgressive genes in introgression from MUS into CAS2 have been fixed in hybrids, suggesting the importance of related biological functions (reproduction and development, metabolism, cell transportation and immune response) to the adaptation of hybrids in southern China. What need to notice is that previous study proposed that, compared with genes functioning at the cell periphery, genes functioning in the cytoplasm and nucleus are not prone to introgression (Janoušek et al. 2015), which is not the case according to our results in all four hybrid groups with whole-genome data.

In summary, our results provide comprehensive insights into the origin, divergence and inter-subspecies hybridization of house mice, which can serve as valuable references for future studies on the evolutionary biology of house mice. One limitation of this work is that available whole-genome data for hybrids from Japan and Europe are limited, which may resulted in the under-estimation of adaptive introgression in these two hybrid groups. In addition, the mechanism for ultra-strong introgression ability of the MUS-type Y chromosome into other subspecies need to explore thoroughly in future.

## Methods

### Re-sequencing, read mapping and SNP calling

A total of 242 wild mice trapped in China were identified using mitochondrial cytochrome oxidase subunit I (COI) as a barcode. Genomic DNA of each sample was extracted from small pieces of tail using a QIAGEN DNA Extraction Kit (Qiagen, USA). The quality and integrity of extracted DNA were assessed with the A260/A280 ratio using a NanoDrop ND-1000 spectrophotometer (Thermo Fisher Scientific Inc., Waltham, MA, USA) and by agarose gel electrophoresis. Libraries with an insert size of ∼300 bp were prepared, and sequencing was performed on an Illumina HiSeq XTen/NovaSeq/BGI platform by a commercial service (Biomarker Technologies, Beijing, China), with 150-bp read lengths. Raw sequencing data sets are deposited in CNCB BioProject under accession number: PRJCA016707. In addition, published raw reads of 157 wild mice (Halligan et al. 2010, accession number: PRJEB2176; Harr et al. 2016, accession number: PRJEB14167, PRJEB11742, PRJEB9450; Fujiwara et al. 2022, accession number: PRJDB11027) were incorporated into the data set, and 399 mouse samples from Eurasia were included in the following analyses. Genomic data for one individual of *Mus spretus* (ERR1124346) published previously were used as an outgroup. Raw sequencing reads were filtered using fastp (v0.20.0) (Chen et al. 2018), and sequencing reads containing adapters and reads with low quality were removed.

The clean reads of autosomes and X chromosomes were aligned to the reference genome GRCm38.p6 (https://www.ncbi.nlm.nih.gov/data-hub/genome/GCF_000001635.26/) using BWA-MEM (v0.7.5a) aligner with default parameters (Li and Durbin 2009). The mapping reads were then converted into BAM files and sorted using SAMtools (v1.10) (Li et al. 2009). Duplicate reads were removed using the MarkDuplicates module in GATK (v4.1.8.0) (McKenna et al. 2010). Raw SNP and insertion/deletion (INDEL) calls were performed using the GATK HaplotypeCaller module with the “-ERC GVCF” option. All genomic variant call format (gVCF) files were merged using the CombinGVCFs module, and variants of all samples were jointly called using the GenotypeGVCFs module, which generated the raw variant call format (VCF) file. The SelectVariants module was utilized to select SNPs and INDELs in the VCF file. To minimize the risk of false-positive calls, quality control of the selected SNPs and INDELs was performed using the VariantFiltering module. The following parameters were applied in SNP filtering: QUAL< 30.0, QD < 2.0, SOR > 3.0, FS > 60.0, MQ < 40.0, MQRankSum < −12.5, ReadPosRankSum < −8.0, --cluster-window-size 10 --cluster-size 3. For INDEL filtering, the applied parameters were QUAL< 30.0, QD < 2.0, FS > 200.0, InbreedingCoeff < −0.8, ReadPosRankSum < −20.0, and SOR > 10.0. After quality control, the multiallelic sites in the data were further removed, and only the biallelic sites were retained.

Kinship inference among samples was performed using KING (Manichaikul et al. 2010) with the “–kinship” option. According to KING, the expected ranges of kinship coefficients were > 0.354 for duplicate/monozygotic twins, > 0.177 and < 0.354 for first-degree relationships, > 0.088 and < 0.177 for second-degree relationships, > 0.044 and < 0.088 for third-degree relationships, and < 0.044 for unrelated individuals. We excluded 50 samples with relationships closer than third-degree. Finally, genomic data of 349 wild samples (209 new samples and 140 published samples) from 19 countries of Eurasia were retained.

The identified SNPs were further annotated by SnpEff (v4.3) (Cingolani et al. 2012). The GRCm38.p6 reference genome and gff3 files were used to manually construct a mouse SnpEff annotation database. According to their locations in the genome, variants were classified into eight groups: intergenic region variant, upstream gene variant, 5 prime UTR variant, intron variant, splice region variant, exon variant, 3 prime UTR variant, and downstream gene variant.

Similarities of deep-sequenced genomes to the GRCm38.p6 reference genome were calculated and visualized as the average of the identity scores of individual SNPs within 1 Mb windows along the genome, according to the formula described previously (Ai et al. 2015).

### Construction of population genetic structure using autosomal and X-chromosomal SNPs

A neighbour-joining phylogenetic tree including 349 individuals was constructed by VCF2Dis (https://github.com/BGI-shenzhen/VCF2Dis) using autosomal and X-chromosomal SNPs. A *P*-distance model was applied in the analyses, and the tree was visualized with ggtree (Yu et al. 2017), with *M. spretus* (SP1) as the outgroup. The population genetic structure was examined using ADMIXTURE (v1.3.0) (Alexander et al. 2009) with *K* values (the putative number of populations) ranging from 1 to 6. The optimal number of populations was selected using cross-validation, and the admixture result was visualized by Pophelper (Francis and Pophelper 2017). Principal component analysis (PCA) was performed using the smartpca program in EIGENSOFT (https://github.com/argriffing/eigensoft), and the VCF file was converted into eigenstrat format by vcf2eigenstrat.py (https://github.com/mathii/gdc). Program output included analysed eigenvectors and eigenvalues, and the first three eigenvectors were selected for graph plotting by ggplot2.

Population relatedness and migration events were inferred using TreeMix (Pickrell and Pritchard 2012). All samples were included in the analyses, with *M. spretus* as the root group. To mitigate the effects of linkage disequilibrium (LD), SNPs were filtered using the “–-indep-pairwise 50 10 0.1” option of PLINK1.9 (Chang et al. 2015), and the VCF file was converted into treemix format by vcf2treemix.sh (https://github.com/speciationgenomics/scripts). TreeMix was applied with the parameters “-noss -bootstrap -k 500 -root SPR” and from 0 to 5 migration events. The R script plotting funcs.R was used to visualize the results, and the best migration events were inferred by residual visualization.

### Estimation of genomic diversity and linkage disequilibrium decay

Population genomic statistics were calculated using VCFtools (v1.9) (Danecek et al. 2011). Ten representatives were chosen from each clade (CAS1, CAS2, DOM and MUS) to calculate nucleotide diversity (π) and fixation index (*F*_ST_) values with a 100 kb window. SNPs with a missing rate <10% (--max-missing 0.9) and minor allele count =1 (--mac 1) were used in the calculation. To estimate and compare the patterns of linkage disequilibrium (LD) among the four clades, the squared correlation coefficients (*r^2^*) between pairwise SNPs were computed using PopLDdecay (v.3.41) (Zhang et al. 2019). The parameters used in the analyses were “--MaxDist 500 --MAF 0.05 --Miss 0.05”. The average *r^2^* value was calculated for pairwise markers in a 500 kb window and averaged across the whole genome. To achieve a smooth curve, Plot_MultiPop.pl was run to plot the LD decay graph with the following parameters: -bin1 100 -bin2 300 -break 300 -maxX 500. To further explore the genetic distance between clades, we used the extract_f2 module in AdmixTools2 (https://github.com/uqrmaie1/admixtools) to calculate the *f*_2_ statistics and then used the *f*_3_ module to calculate the outgroup (SP1) *f*_3_ statistics based on the *f*_2_ statistics.

### Estimation of divergence times of subspecies

The divergence times of three subspecies (DOM, MUS and CAS2) from the ancestral population were inferred using the flexible and robust simulation-based composite-likelihood approach implemented in the fastsimcoal2 program (Excofffier et al. 2021), which infers demographic parameters from the site frequency spectrum (SFS). Six representatives with low rates of missing data for each clade were used in the analyses. Folded SFSs (MAF) were calculated by easySFS (https://github.com/isaacovercast/easySFS). Estimates were obtained from 200,000 simulations per likelihood estimation (-n200,000) using 50 expectation/conditional maximization (ECM) cycles (-L50). Monomorphic sites in the observed SFSs were not considered for parameter inference (−0), and the output comprised all SNPs in the DNA sequences (-s0). The program was run 100 times, and the 95% confidence interval (CI) and point estimates for each fastsimcoal2 parameter were obtained using 1,000 bootstrap replicates in R.

### Simulation of the demographic histories

The population dynamics of CAS1, CAS2, DOM and MUS were simulated using pairwise sequentially Markovian coalescent (PSMC, https://github.com/lh3/psmc) (Li and Durbin 2011) with deep-sequenced genomes (coverage depth ≥ 18×) (Nadachowska-Brzyska et al. 2016). The psmc function in the PSMC package was run to estimate historical effective population size (*Ne*) with the following parameters: -N25 -t15 -r5 -p ‘4+25+2+4+6’, 1.5 generations per year (Phifer-Rixey et al. 2020), and a mutation rate of 4.1 × 10^−9^ per site per generation (Geraldes et al. 2008). To perform bootstrapping, the splitfa function in the PSMC package was run to split long chromosome sequences into shorter segments. When the ‘-b’ option is applied, psmc will randomly sample with replacement from these segments. For each clade, 9 to 20 individuals from different geographic populations were included in the analyses, and 100 bootstrap estimates were generated for representative individuals of each clade (CAS1: PA1; CAS2: ID2; DOM: GER2 and IR2; MUS: KA5).

Recent population dynamics of MUS and CAS2 were inferred using SMC^++^ (https://github.com/popgenmethods/smcpp) (Terhorst et al. 2017). Four individuals of MUS from Xinjiang, China, and four samples of CAS2 from the south coast of China were selected, and the related VCF file was converted into the SMC^++^ input format using vcf2smc tools. SMC^++^ inferences were performed for various sets of genotypes using the SMC^++^ estimate command with the following parameters: --spline cubic -knots 15 - timepoints 100 10000000 -cores 20. Generations were calculated with a mutation rate of 4.1 × 10^−9^ per site per generation (Geraldes et al. 2008) and 1.5 generations per year (Phifer-Rixey et al. 2020). The SMC^++^ plot command was run to plot raw data with the ‘-c’ option and then visualized in R using the plot function. The effective population size (*Ne*) over the past 100 years (CAS2: 15,000, MUS: 35,000) was used to estimate recombination rates along the genomes of MUS and CAS2 in subsequent analyses.

### Construction of paternal phylogeny

High-throughput sequencing reads of all 198 males were mapped to the Y-chromosome reference sequence in GRCm38.p6 using BWA-MEM aligner. Raw SNP calling and filtering were performed using GATK. To eliminate the influence of the X chromosome, we excluded SNPs called from the VCF file for female individuals whose reads were misaligned with the Y-chromosome reference sequence. Heterozygous sites were also removed, and only sites with missing rates less than 20% were retained. After filtering, 10,874 high-quality SNPs were obtained and used to construct the neighbour-joining phylogenetic tree using VCF2Dis.

BCFtools (v1.10.2) was applied to select SNPs located in ancestral single-copy regions (Soh et al. 2014) on the short arm of the mouse Y chromosome. It was ensured that the minor allele frequency at each site was greater than 0.05. Total of 546 SNPs were screened out, and the VCF file was then converted into a NEXUS haplotype data file by vcf2phylip.py (https://github.com/edgardomortiz/vcf2phylip) and DnaSP6 (Rozas et al. 2017). The statistical parsimony networks (Clement et al. 2000) were constructed using PopART^61^(Leigh and Bryant 2015).

### Evaluation on genomic introgression between different clades with *D*-statistics

*D*-statistics^62^ (Durand et al. 2011) based on ABBA and BABA SNP frequency differences were applied to detect potential gene flow between different clades of house mice. A tree topology from sites covered by the four populations in the form [[[H1, H2], H3], O] was used to test whether two conspecific populations (H1 and H2) shared more alleles with a candidate donor, H3. *M. spretus* (SP1) was used as the outgroup (O). For sites where SP1 possessed the ancestral allele “A” and H3 had the derived allele “B”, we calculated the frequency of the “ABBA” pattern versus the “BABA” pattern in the topology. The candidate recipient was defined as H2. The “Dtrios module” of Dsuite (Malinsky et al. 2021) was used to calculate the *D* value and *f*_4_ ratio for all populations. The *D* value and Z score (|Z score| > 3) (Chen et al. 2021) were used to indicate the mean degree of introgression in each population. The *f*_4_ ratio was used to estimate the admixture proportions in the recipient. The calculation of the *f*_4_ admixture ratio required H3 to be split into two subsets (H3a and H3b) by randomly sampling alleles from H3 at each SNP.

*D*-statistics based on BABA and ABBA SNP frequency differences were applied to detect potential hybrids between two clades of house mice. We used the *f*_4_ module of AdmixTools2 with the ‘f4mode = FALSE’ option, enabling calculation of the *D* value. In the [[[H1, H2], H3], O] tree topology with four individuals, the candidate recipient and donor were defined as H1 and H3, respectively. The calculation was based on “BABA - ABBA” statistics. *M. spretus* (SP1) was used as the outgroup (O). The possibilities of introgression between MUS and CAS2 in China, between CAS1 and CAS2 on the Indian subcontinent, in Southeast Asia and in southern China, between CAS1 and MUS in Afghanistan, between CAS2 and DOM in Southeast Asia and southern China, and between DOM and MUS in Central Asia, Iran and Europe were analysed. The statistical significance of the *D*-value was evaluated using a Z test, and H1 individuals with *D* > 0.05 and Z > 3 were considered hybrids between two clades.

### Identifying introgressive genomic regions with ABBA-BABA statistics

For phasing genotypes and filling missing genotypes, Beagle (v5.2) (Browning et al. 2021) was used to process our VCF file. To identify introgressive genomic regions, hybrids with introgression from MUS into CAS2 in China, hybrids with introgression from CAS2 into MUS in China, hybrids with introgression from CAS2 into MUS in Japan, and hybrids with introgression from DOM into MUS in the Czech Republic were merged into one hybrid group. ParseVCF.py was performed to convert the VCF format into the input format of ABBABABAwindows.py. Then, the ABBABABAwindows.py script (https://github.com/simonhmartin/genomics_general) was used to calculate the *D* statistic, *f*_d_ (Martin et al. 2015) and *f*_dM_ (Malinsky et al. 2015) in sliding windows across the genome with the parameters “-f phased -w 100000 -s 20000 -m 100 -T 10”. *f*_d_ is often used to localize genomic regions with significant introgression, and *f*_dM_ is an alternative statistic of *f*_d_ used in this case to analyse shared variation between H3 and H2 (positive *D*) or between H3 and H1 (negative *D*). The popgenWindows.py script (https://github.com/simonhmartin/genomics_general) was used to calculate standard population genomic statistics (π, *F*_ST_, *d*_xy_) in sliding windows using the parameters “-f phased -w 100000 -s 20000 -m 100 -T 10”. Manhattan plots were generated with the R package CMplot (https://github.com/YinLiLin/R-CMplot). For hybrids in China and Japan, the top 5% values of the empirical genome-wide distribution of *f*_dM_ were considered introgressive regions due to the excessive degree of introgression. For hybrids in the Czech Republic, genomic regions with FDR values < 0.05 were considered introgressive regions due to the lower level of introgression. Then, the adjacent introgressive windows were merged by an in-house R script, and the lengths of introgressive segments were measured. Introgressive regions containing high-*F*_ST_ outliers (corresponding to the top 5% values of the empirical genome-wide distribution of *F*_ST_) and high-π ratio outliers (corresponding to the top 5% values of the empirical genome-wide distribution of the π ratio) were considered regions of adaptive introgression. Introgression on the X chromosome between CAS2 and MUS was analysed separately using the same method.

### Estimation of local recombination rates along the genomes of CAS2 and MUS

A machine learning approach implemented in FastEPRR (v2.0) (Gao et al. 2016) with a 100 kb window was applied to estimate the recombination rates of various regions across the genomes of CAS2 and MUS. In the FastEPRR results (*Rho* = 4*Ner*), *Rho* is the frequency-weighted chromosomal recombination rate, *Ne* is the effective population size, and *r* is the recombination rate of the window. The mean local recombination rates along the genomes were calculated with the above formula, where *Ne* applied the results of SMC^++^ evaluation over the past 100 years.

### Detection of incomplete lineage sorting

To distinguish genomic introgressive regions from the influence of incomplete lineage sorting (ILS), we calculated the probability of the alternative scenario of ILS. The expected length of a shared ancestral sequence is *L* = 1/(*r*×*t*) (Huerta-Sánchez et al. 2014), where “*r*” is the recombination rate per generation per base pair and “*t*” is the branch length between MUS and CAS2 since their divergence. Using the length of each introgressive tract as the expected length and the inferred recombination rates of local regions, we estimated the possible ages of each introgressive region (*t* = 1/(*L*×*r*)). The *P* values were calculated according to Z values using the standard normal distribution and then further corrected for multiple testing by using the BenjaminiDHochberg false discovery rate (FDR) method. FDR values < 0.05 were considered outliers. The probability of a fragment with length > *m* is 1 - GammaCDF (*m*, shape = 2, rate = 1/*L*), where GammaCDF is the cumulative distribution function (CDF) of the gamma distribution and “m” is the length of introgressive segments. The gamma distribution was implemented through the pgamma function in R.

### Gene identification and functional enrichment analyses

Genes in the target regions were identified by bedtools (v2.27.1) (Quinlan and Hall 2010) with a GRCm38.p6 gtf file (https://www.ncbi.nlm.nih.gov/datahub/genome/GCF_000001635.26/). The “bedtools intersect” command was used to merge the overlapping target windows and obtain the target genes. Gene Ontology (GO) and Kyoto Encyclopedia of Genes and Genomes (KEGG) analyses were performed with the clusterProfiler toolkit (v4.4.4) (Yu et al. 2012) and org.Mm.eg.db package (v3.15.0) in R to obtain functional classifications and enrichment pathways of candidate genes, respectively. The enrichment analyses were performed using all *Mus musculus* gene annotations as background. The maximum number of genes for the enriched term was set to 2,000 with the ‘maxGSSize’ option. The Benjamini-Hochberg FDR method was used to correct the resulting *P* values. GO annotations with adjusted *P* < 0.05 and pathways with *P* < 0.05 were considered significantly enriched.

## Data access

All raw and processed sequencing data generated in this study have been submitted to the CNCB BioProject database (https://www.cncb.ac.cn/) under accession number PRJCA016707.

## Code availability

All computational analyses are publicly available on GitHub (https://github.com/chen1238/Populations-Genetic-Analysis).

## Acknowledgements

This research was made possible by grants from the National Natural Science Foundation of China to L. H. (32070403) and M. J. (32370447) and from the Second Tibetan Plateau Scientific Expedition and Research Program to M. J. (2019QZKK05010303). H. Yu at National Taiwan University kindly provided mice samples from the Taiwan island used in this study.

## References

Ai H, Fang X, Yang B, Huang Z, Chen H, Mao L, Zhang F, Zhang L, Cui L, He W, et al. 2015. Adaptation and possible ancient interspecies introgression in pigs identified by whole-genome sequencing. Nat Genet 47: 217–225. doi: 10.1038/ng.3199

Alexander DH, Novembre J, Lange K. 2009. Fast model-based estimation of ancestry in unrelated individuals. Genome Res 19: 1655–1664. doi: 10.1101/gr.094052.109

Bonhomme F, Orth A, Cucchi T, Rajabi-Maham H, Catalan J, Boursot P, Auffray JC, Britton-Davidian J. 2011. Genetic differentiation of the house mouse around the Mediterranean basin: matrilineal footprints of early and late colonization. Proc Biol Sci 278: 1034–1043. doi: 10.1098/rspb.2010.1228

Bonhomme F, Searle JB. 2012. House mouse phylogeography. In Evolution of the House Mouse, (ed. Macholan M, et al.), pp 278–296. Cambridge University Press, Cambridge.

Boursot P, Auffray JC, Britton-Davidian J, Bonhomme F. 1993. The evolution of house mice. Annu Rev Ecol Syst 24: 119–152.

Boursot P, Din W, Anand R, Darviche D, Dod B, Von Deimling F, Talwar GP, Bonhomme F. 1996. Origin and radiation of the house mouse: mitochondrial DNA phylogeny. J Evolution Biol 9: 391–415.

Browning BL, Tian X, Zhou Y, Browning SR. 2021. Fast two-stage phasing of large-scale sequence data. Am J Hum Genet 108: 1880–1890. doi: 10.1016/j.ajhg.2021.08.005

Campbell P, Nachman MW. 2014. X-y interactions underlie sperm head abnormality in hybrid male house mice. Genetics 196: 1231–1240. doi: 10.1534/genetics.114.161703

Chang CC, Chow CC,Tellier LC, Vattikuti S, Purcell SM, Lee JJ. 2015. Second-generation PLINK: rising to the challenge of larger and richer datasets. Gigascience 4: s13742-015-0047-8. doi: 10.1186/s13742-015-0047-8

Chen S, Zhou Y, Chen Y, Gu J. 2018. fastp: an ultra-fast all-in-one FASTQ preprocessor. Bioinformatics 34: i884–i890. doi: 10.1093/bioinformatics/bty560

Chen T, Li J, Xu M, Zhao Q, Hou Y, Yao L, Zhong Y, Chou P, Zhang W, Zhou P, et al. 2016. PKCε phosphorylates MIIP and promotes colorectal cancer metastasis through inhibition of RelA deacetylation. Nat Commun 8: 939. doi: 10.1038/s41467-017-01024-2

Chen N, Cai Y, Chen Q, Li R, Wang K, Huang Y, Hu S, Huang S, Zhang H, Zheng Z, et al. 2018. Whole-genome resequencing reveals world-wide ancestry and adaptive introgression events of domesticated cattle in East Asia. Nat Commun 9: 2337.doi: 10.1038/s41467-018-04737-0

Chen Z, Xu Y, Xie X, Wang D, Aguilar-Gómez D, Liu G, Li X, Esmailizadeh A, Rezaei V, Kantanen J, Ammosov I, et al. 2021. Whole-genome sequence analysis unveils different origins of European and Asiatic mouflon and domestication-related genes in sheep. Commun Biol 4: 1307. doi: 10.1038/s42003-021-02817-4

Christmas MJ, Jones JC, Olsson A, Wallerman O, Bunikis I, Kierczak M, Peona V, Whitley KM, Larva T, Suh A, et al. Genetic barriers to historical gene flow between cryptic species of Alpine bumblebees revealed by comparative population genomics. Mol Biol Evol 38: 3126–3143. doi: 10.1093/molbev/msab086

Cingolani P, Platts A, Wang LL, Coon M, Nguyen T, Wang L, Land SJ, Lu X, Ruden DM. 2012. A program for annotating and predicting the effects of single nucleotide polymorphisms, SnpEff: SNPs in the genome of *Drosophila melanogaster* strain w1118; iso-2; iso-3. Fly (Austin) 6: 80–92. doi: 10.4161/fly.19695

Clement M, Posada D, Crandall KA. 2000. TCS: a computer program to estimate gene genealogies. Mol Ecol 9: 1657–1659. doi: 10.1046/j.1365-294x.2000.01020.x

Danecek P, Auton A, Abecasis G, Albers CA, Banks E, DePristo MA, Handsaker RE, Lunter G, Marth GT, Sherry ST, et al. 2011. The variant call format and VCFtools. Bioinformatics 27: 2156–2158. doi: 10.1093/bioinformatics/btr330

Del Punta K, Leinders-Zufall T, Rodriguez I, Jukam D, Wysocki CJ, Ogawa S, Zufall F, Mombaerts P. 2002. Deficient pheromone responses in mice lacking a cluster of vomeronasal receptor genes. Nature 419: 70–74. doi: 10.1038/nature00955

Dod B, Jermiin LS, Boursot P, Chapman VH, Nielsen JT, Bonhomme F. 1993. Counterselection on sex chromosomes in the *Mus musculus* European hybrid zone. J Evol Biol 6: 529–546.

Doran J, Walters C, Kyle V, Wooding P, Hammett-Burke R, Colledge WH. 2016. *Mfsd14a* (*Hiat1*) gene disruption causes globozoospermia and infertility in male mice. Reproduction 152: 91–99. doi: 10.1530/REP-15-0557

Durand EY, Patterson N, Reich D, Slatkin M. 2011. Testing for ancient admixture between closely related populations. Mol Biol Evol 28: 2239–2252.

Duvaux L, Belkhir K, Boulesteix M, Boursot P. 2011. Isolation and gene flow: inferring the speciation history of European house mice. Mol Ecol 20: 5248–5264. doi: 10.1111/j.1365-294X.2011.05343.x

Excofffier L, Marchi N, Marques DA, Matthey-Doret R, Gouy A, Sousa VC. 2021. fastsimcoal2: demographic inference under complex evolutionary scenarios. Bioinformatics 37: 4882–4885. doi: 10.1093/bioinformatics/btab468

Feng C, Wang J, Liston A, Kang M. 2023. Recombination variation shapes phylogeny and introgression in wild diploid strawberries. Mol Biol Evol 40: msad049. doi: 10.1093/molbev/msad049

Francis RM. 2017. Pophelper: an R package and web app to analyze and visualize population structure. Mol Ecol Resour 17: 27–32. doi:10.1111/1755-0998.12509

Fu R, Zhu Y, Liu Y, Feng Y, Lu RS, Li Y, Li P, Kremer A, Lascoux M, Chen J. 2022. Genome-wide analyses of introgression between two sympatric Asian oak species. Nat Ecol Evol 6: 924–935. doi: 10.1038/s41559-022-01754-7

Fujiwara, K. Kawai Y, Takada T, Shiroishi T, Saitou N, Suzuki H, Osada N. 2022. Insights into *Mus musculus* population structure across Eurasia revealed by whole-genome analysis. Genome Biol Evol 14: evac068. doi: 10.1093/gbe/evac068

Gao F, Ming C, Hu W, Li H. 2016. New software for the fast estimation of population recombination rates (FastEPRR) in the genomic era. G3 (Bethesda) 6: 1563–1571. doi: 10.1534/g3.116.028233

Geraldes A, Basset P, Gibson B, Smith KL, Harr B, Yu HT, Bulatova N, Ziv Y, Nachman MW. 2008. Inferring the history of speciation in house mice from autosomal, X-linked, Y-linked and mitochondrial genes. Mol Ecol 17(24): 5349–5363. doi: 10.1111/j.1365-294X.2008.04005.x

Geraldes A, Basset P, Smith KL, Nachman MW. 2011. Higher differentiation among subspecies of the house mouse (*Mus musculus*) in genomic regions with low recombination. Mol Ecol 20: 4722–4736. doi: 10.1111/j.1365-294X.2011.05285.x

Halligan DL, Oliver F, Eyre-Walker A, Harr B, Keightley PD. 2010. Evidence for pervasive adaptive protein evolution in wild mice. PLoS Genet 6: e1000825. doi: 10.1371/journal.pgen.1000825

Hardouin EA, Orth A, Teschke Meike, Darvish J, Tautz D, Bonhomme F. 2015. Eurasian house mouse (*Mus musculus* L.) differentiation at microsatellite loci identifies the Iranian plateau as a phylogeographic hotspot. BMC Evol Biol 15: 26. doi: 10.1186/s12862-015-0306-4

Harr, B. Karakoc E, Neme R, Teschke M, Pfeifle C, Pezer Ž, Babiker H, Linnenbrink M, Montero I, Scavetta R, et al. 2016. Genomic resources for wild populations of the house mouse, *Mus musculus* and its close relative *Mus spretus*. Sci Data 3: 160075. doi: 10.1038/sdata.2016.75

Hewitt, G. 2000. The genetic legacy of the Quaternary ice ages. Nature 405: 907–913. doi: 10.1038/35016000

Huerta-Sánchez E, Jin X, Asan, Bianba Z, Peter BM, Vinckenbosch N, Liang Y, Yi X, He M, Somel M, et al. 2014. Altitude adaptation in Tibetans caused by introgression of Denisovan-like DNA. Nature 512: 194–197. doi: 10.1038/nature13408

Jacobs S, Ruusuvuori E, Sipilä ST, Haapanen A, Damkier HH, Kurth I, Hentschke M, Schweizer M, Rudhard Y, Laatikainen LM, et al. 2008. Mice with targeted *Slc4a10* gene disruption have small brain ventricles and show reduced neuronal excitability. Proc Natl Acad Sci USA 105: 311–316. doi: 10.1073/pnas.0705487105

Janoušek V, Munclinger P, Wang L, Teeter KC, Tucker PK. 2015. Functional organization of the genome may shape the species boundary in the house mouse. Mol Biol Evol 32: 1208–1220. doi: 10.1093/molbev/msv011

Jing M, Yu H, Bi X, Lai Y, Jiang W, Huang L. 2014. Phylogeography of Chinese house mice (*Mus musculus musculus*/*castaneus*): distribution, routes of colonization and geographic regions of hybridization. Mol Ecol 23: 4387–4405. doi: 10.1111/mec.12873

Jones EP, Van der Kooij J, Solheim R, Searle JB. 2010. Norwegian house mice (*Mus musculus musculus*/*domesticus*): distributions, routes of colonization and patterns of hybridization. Mol Ecol 19: 5252–5264. doi: 10.1111/j.1365-294X.2010.04874.x

Jones MK, Liu X. 2009. Origins of agriculture in East Asia. Science 324: 730–731. doi: 10.1126/science.1172082

Leigh JW, Bryant D. 2015. POPART: Full-feature software for haplotype network construction. Meth Ecol Evol 6: 1110–1116.doi: 10.1111/2041-210X.12410

Li H, Durbin R. 2009. Fast and accurate short read alignment with Burrows-Wheeler transform. Bioinformatics 25: 1754–1760. doi: 10.1093/bioinformatics/btp324

Li H, Durbin R. 2011. Inference of human population history from individual whole-genome sequences. Nature 475: 493–496. doi: 10.1038/nature10231

Li, H. Handsaker B, Wysoker A, Fennell T, Ruan J, Homer N, Marth G, Abecasis G, Durbin Richard,1000 Genome Project Data Processing Subgroup. 2009. The Sequence Alignment/Map format and SAMtools. Bioinformatics 25: 2078–2079. doi: 10.1093/bioinformatics/btp352

Liu Y, Takahashi A, Kitano T, Koide T, Shiroishi T, Moriwaki K, Saitou N. 2008.Mosaic genealogy of the *Mus musculus* genome revealed by 21 nuclear genes from its three subspecies. Genes Genet Syst 83: 77–88. doi: 10.1266/ggs.83.77

Lundrigan B, Jansa S, Tucker P. 2002. Phylogenetic relationships in the genus *Mus*, based on paternally, maternally, and biparentally inherited characters. Syst Biol 51: 410–431. doi: 10.1080/10635150290069878

Macholan M, Munclinger P, Sugerková M, Dufková P, Bímová B, Bozíková E, Zima J, Piálek J. 2007. Genetic analysis of autosomal and X-linked markers across a mouse hybrid zone. Evolution 61: 746–771. doi: 10.1111/j.1558-5646.2007.00065.x

Malinsky M, Challis RJ, Tyers AM, Schiffels S, Terai Y, Ngatunga BP, Miska EA, Durbin R, Genner MJ, Turne GF. 2015. Genomic islands of speciation separate cichlid ecomorphs in an East African crater lake. Science 350, 1493–1498. doi: 10.1126/science.aac9927

Malinsky M, Matschiner M, Svardal H. 2021. Dsuite - Fast D-statistics and related admixture evidence from VCF files. Mol Ecol Resour 21: 584–595. doi: 10.1111/1755-0998.13265

Manichaikul A, Mychaleckyj JC, Rich SS, Daly K, Sale M, Chen W. 2010. Robust relationship inference in genome-wide association studies. Bioinformatics 26: 2867–2873. doi: 10.1093/bioinformatics/btq559

Martin SH, Davey JW, Jiggins CD. 2015. Evaluating the use of ABBA-BABA statistics to locate introgressed loci. Mol Biol Evol32: 244–257. doi: 10.1093/molbev/msu269

Martin SH, Davey JW, Salazar C, Jiggins CD. 2019. Recombination rate variation shapes barriers to introgression across butterfly genomes. PLoS Biol 17: e2006288. doi: 10.1371/journal.pbio.2006288

McKenna A, Hanna M, Banks E, Sivachenko A, Cibulskis K, Kernytsky A, Garimella K, Altshuler D, Gabriel S, Daly M, et al. 2010. The Genome Analysis Toolkit: a MapReduce framework for analyzing next-generation DNA sequencing data. Genome Res. 20, 1297–1303. doi: 10.1101/gr.107524.110

Morgan AP, Pardo-Manuel de Villena F. 2017. Sequence and structural diversity of mouse Y chromosomes. MolBiol Evol 34: 3186–3204. doi: 10.1093/molbev/msx250

Munclinger P, Božíková E, Šugerková M, Piálek J, Macholán M. 2002. Genetic variation in house mice (*Mus*, Muridae, Rodentia) from the Czech and Slovak Republics. Folia Zool 51: 81–92.

Nadachowska-Brzyska K, Burri R, Smeds L, Ellegren H. 2016. PSMC analysis of effective population sizes in molecular ecology and its application to black-and-white *Ficedula* flycatchers. Mol Ecol 25: 1058–1072. doi: 10.1111/mec.13540

Payseur BA, Krenz JG,Nachman MW. Differential patterns of introgression across the X chromosome in a hybrid zone between two species of house mice. Evolution 58(9), 2064–2078 (2004). doi: 10.1111/j.0014-3820.2004.tb00490.x

Phifer-Rixey M, Harr B, Hey J. 2020. Further resolution of the house mouse (*Mus musculus*) phylogeny by integration over isolation-with-migration histories. BMC Evol Bio 20: 120. doi: 10.1186/s12862-020-01666-9

Phifer-Rixey M, Nachman MW. 2015. Insights into mammalian biology from the wild house mouse *Mus musculus*. ELife 4: e05959. doi: 10.7554/eLife.05959

Pickrell JK, Pritchard JK. 2012. Inference of population splits and mixtures from genome-wide allele frequency data. PLoS Genet. 8: e1002967. doi: 10.1371/journal.pgen.1002967

Potter PK, Bowl MR, Jeyarajan P, Wisby L, Blease A, Goldsworthy ME, Simon MM, Greenaway S, Michel V, Barnard A, et al. 2016. Novel gene function revealed by mouse mutagenesis screens for models of agerelated disease. Nat Commun 7: 12444. doi: 10.1038/ncomms12444

Quinlan AR, HallI M. 2010. BEDTools: a flexible suite of utilities for comparing genomic features. Bioinformatics 26: 841–842. doi: 10.1093/bioinformatics/btq033

Rajabi-Maham H, Orth A, Bonhomme F. 2008. Phylogeography and postglacial expansion of *Mus musculus domesticus* inferred from mitochondrial DNA coalescent, from Iran to Europe. Mol Ecol 17: 627–641. doi: 10.1111/j.1365-294X.2007.03601.x

Rozas J, Ferrer-Mata A, Sánchez-DelBarrio JC, Guirao-Rico S, Librado P, Ramos-Onsins SE, Sánchez-Gracia A. 2017. DnaSP 6: DNA Sequence Polymorphism Analysis of Large Data Sets. Mol Biol Evol 34: 3299–3302. doi: 10.1093/molbev/msx248

Searle JB, Jones CS, Gündüz I, Scascitelli M, Jones EP, Herman JS, Rambau RV, Noble LR, Berry RJ, Giménez MD, et al. 2009. Of mice and (Viking?) men: phylogeography of British and Irish house mice. Proc Biol Sci 276: 201–207. doi: 10.1098/rspb.2008.0958

Soh YQS, Alföldi J, Pyntikova T, Brown LG, Graves T, Minx PJ, Fulton RS, Kremitzki C, Koutseva N, Mueller JL, et al. 2014. Sequencing the mouse Y chromosome reveals convergent gene acquisition and amplification on both sex chromosomes. Cell 159: 800–813. doi: 10.1016/j.cell.2014.09.052

Teeter KC, Payseur B A, Harris LW, Bakewell MA, Thibodeau LM, O’Brien JE, Krenz JG, Sans-Fuentes MA, Nachman MW, Tucker PK. 2008. Genome-wide patterns of gene flow across a house mouse hybrid zone. Genome Res 18: 67–76. doi: 10.1101/gr.6757907

Teeter KC, Thibodeau LM, Gompert Z, Buerkle CA, Nachman MW, Tucker PK. 2010. The variable genomic architecture of isolation between hybridizing species of house mice. Evolution 64: 472–485. doi: 10.1111/j.1558-5646.2009.00846.x

Teng H, Zhang Y, Shi C, Mao F, Cai W, Lu L, Zhao F, Sun Z, Zhang J. 2017. Population genomics reveals speciation and introgression between brown Norway rats and their sibling species. Mol Biol Evol 34: 2214–2228. doi: 10.1093/molbev/msx157

Terashima M, Furusawa S, Hanzawa N, Tsuchiya K, Suyanto A, Moriwaki K, Yonekawa H, Suzuki H. 2006. Phylogeographic origin of Hokkaido house mice (*Mus musculus*) as indicated by genetic markers with maternal, paternal and biparental inheritance. Heredity 96, 128–138. doi: 10.1038/sj.hdy.6800761

Terhorst J, Kamm JA, Song YS. 2017. Robust and scalable inference of population history from hundreds of unphased whole genomes. Nat Genet 49: 303–309. doi: 10.1038/ng.3748

Turner LM, Schwahn DJ, Harr B. 2012. Reduced male fertility is common but highly variable in form and severity in a nature house mouse hybrid zone. Evolution 66: 443–458. doi: 10.1111/j.1558-5646.2011.01445.x

Ullrich KK, Tautz D. 2020. Population genomics of the house mouse and the brown rat. In Statistical population genomics (ed. Dutheil JY), pp. 435–452. Humana Press, New Jersey.

Wang T, Xiao Y, Hu Z, Gu J, Hua R, Hai Z, Chen X, Zhang J, Yu Z, Wu T, et al. 2022. MFN2 deficiency impairs mitochondrial functions and PPAR pathway during spermatogenesis and meiosis in mice. Front Cell Dev Biol 10: 862506. doi: 10.3389/fcell.2022.862506

Wang Z, Shi Y, Ying C, Jiang Y, Hu J. 2021. Hypoxia-induced PLOD1 overexpression contributes to the malignant phenotype of glioblastoma via NF-κB signaling. Oncogene 40: 1458–1475. doi: 10.1038/s41388-020-01635-y

Xin A, Qu R, Chen G, Zhang L, Chen J, Tao C, Fu J, Tang J, Ru Y, Chen Y, et al. 2020. Disruption in ACTL7A causes acrosomal ultrastructural defects in human and mouse sperm as a novel male factor inducing early embryonic arrest. Sci Adv 6: eaaz4796. doi: 10.1126/sciadv.aaz4796

Yap MW, Colbeck E, Ellis SA, Stoye JP. 2014. Evolution of the retroviral restriction gene Fv1: inhibition of non-MLV retroviruses. PLoS Pathog 10: e1003968. doi: 10.1371/journal.ppat.1003968

Yi C, Zhu Z, Wei L, Cui Z, Zheng B, Shi Y. 2007. Advances in numerical dating of quaternary glaciations in China. Z Geomorphol 51 (Sup 2): 153–175. doi: 10.1127/0372-8854/2007/0051S2-0153

Yonekawa H, Moriwaki K, Gotoh O, Miyashita N, Matsushima Y, Shi LM, Cho WS, Zhen XL, Tagashira Y. 1988. Hybrid origin of Japanese mice ‘*Mus musculus molossinus*’: evidence from restriction analysis of mitochondrial DNA. MolBiol Evol 5: 63–78. doi: 10.1093/oxfordjournals.molbev.a040476

Yu G, Smith DK, Zhu H, Guan Y, Lam T Y. 2017. ggtree: an r package for visualization and annotation of phylogenetic trees with their covariates and other associated data. Methods Ecol Evol 8: 28–36. doi:10.1111/2041-210X.12628

Yu G, Wang L, Han Y, He Q. 2012. clusterProfiler: an R package for comparing biological themes among gene clusters. Omics 16: 284–287. doi: 10.1089/omi.2011.0118

Zhang C, Dong S, Xu J, He W,Yang T. 2019. PopLDdecay: a fast and effective tool for linkage disequilibrium decay analysis based on variant call format files. Bioinformatics 35: 1786–1788. doi: 10.1093/bioinformatics/bty875

Zheng Z, Wang X, Li M, Li Y, Yang Z, Wang X, Pan X, Gong M, Zhang Y, Guo Y, et al. 2020. The origin of domestication genes in goats. Sci. Adv. 6, eaaz5216. doi: 10.1126/sciadv.aaz5216

